# Transcriptional dysregulation study reveals a core network involving the genesis for Alzheimer’s disease

**DOI:** 10.1101/240002

**Authors:** Guofeng Meng, Hongkang Mei

## Abstract

**Background:** The pathogenesis of Alzheimer’s disease is associated with dysregulation at different levels from transcriptome to cellular functioning. Such complexity necessitates investigations of disease etiology to be carried out considering multiple aspects of the disease and the use of independent strategies. The established works more emphasized on the structural organization of gene regulatory network while neglecting the internal regulation changes.

**Methods:** Applying a strategy different from popularly used co-expression network analysis, this study investigated the transcriptional dysregulations during the transition from normal to disease states.

**Results:** 97 genes were predicted as dysregulated genes, which were also associated with clinical outcomes of Alzheimer’s disease. Both the co-expression and differential co-expression analysis suggested these genes to be interconnected as a core network and that their regulations were strengthened during the transition to disease states. Functional studies suggested the dysregulated genes to be associated with aging and synaptic function. Further, we checked the evolutionary conservation of the gene co-expression and found that human and mouse brain might have divergent transcriptional co-regulation even when they had conserved gene expression profiles.

**Conclusion:** Overall, our study reveals a profile of transcriptional dysregulation in the genesis of Alzheimer’s disease by forming a core network with altered regulation; the core network is associated with Alzheimer’s diseases by affecting the aging and synaptic functions related genes; the gene regulation in brain may not be conservative between human and mouse.

## Background

Alzheimer’s disease (AD) is a neurodegenerative disorder most prevalent in people over the age of 65 years [1, 2, 3]. As the population ages, AD will impact more people and place an increasing economic burden on society [4, 5]. There is still no effective treatment that prevents or slows the disease progression. A significant challenge is the poor understanding of the etiology of this disease [6, 7, 8]. The progression of AD is associated with the dysregulation of many genes at different regulatory levels from transcriptome to neuronal function [9, 10]. To study such a complex multifactorial disease, integrated and large-scale data are necessary to catch the diverse regulatory interactions [11, 12]. The availability of high-throughput transcriptomic sequencing data and clinical annotation in the `Accelerating Medicines Partnership-Alzheimer’s Disease (AMP-AD)` program (provides an opportunity to study the altered transcriptional regulation during the genesis of AD [13, 14, 15, 12].

Co-expression network analysis is useful to infer causal mechanisms for complex diseases [16, 17, 18]. It is based on the assumption that co-expressed genes are usually regulated by the same transcriptional regulators, pathways or protein complexes and that the co-regulated genes can be revealed by analysis of the topological structure of co-expression networks [19, 20]. The co-regulated clusters in the network provide the chance to track the affected pathways or biological processes in diseases [21]. One of the most popular strategies is to find the topological structural changes of the co-expression network under different disease states where these changes indicate regulatory dysregulation in the disease [18, 22]. Another way is to associate the subnetwork expression to disease progresses or clinical traits, which can uncover the regulatory components involved in the dysregulation of diseases [23].

Application of co-expression network analysis to AD data has revealed many AD associated genes and pathways [12]. However, the nature of such analyses may bias toward the genes with higher connectivity in a co-expression network. The complexity of AD genesis leads us to extend co-expression analysis to a more detailed investigation of co-expressed genes, especially for the genes with relatively low connectivity, during the disease progression. To understand the etiology of AD, the changes of regulation are supposed to be more essential than the the regulations themselves. We studied the transcriptional dysregulation by evaluating all the gene pair combinations for their co-expression changes, which can be referred as differential co-expression (DCE) analysis. The genes with altered co-expression are indicated in the genesis of AD and their importance can be ranked by the numbers of changed connections.

In this study, we collected the RNA-seq expression data for 1667 human brain samples from the AMP-AD program. Differential co-expression analysis indicated 87,539 gene pairs to have significant co-expression changes. Among them, 97 genes, including 10 transcription factors, were found to be dysregulated in AD genesis. Both the co-expression and differential co-expression analysis suggested these genes to take roles as a interconnected core network. In the transition from normal to disease states, the co-expression is strengthened in this network. Functional studies supported this network to be involved in the etiology of AD by directly or indirectly affecting aging, synaptic function and metabolism related genes. We also evaluated their evolutionary conservation in mouse. Although the genomic gene expression profiles are conserved, the co-expression patterns were not in mouse, including the core network, which may indicate transcriptional regulation divergence between human and mouse.

## Methods and Materials

### Data collection, processing and quality control

The human expression data were collected from `Accelerating Medicines Partnership-Alzheimer’s Disease (AMP-AD`, http://www.synapse.org/#!Synapse:syn2580853/wiki/66722) program compiling with the data access control at http://dx.doi.Org/doi:10.7303/syn2580853. The RNA-seq expression data from four projects were used, including (1) ROSMAP; (2) MSBB (3) MayoPilot and (4) MayoBB. Based on the clinical annotation, the samples were grouped as AD and normal samples. In this step, some patients with vague disease status, missing annotation or other disease annotations were filtered out.

For the RNA-seq data from each project, the AD and normal samples were separately processed for quality control. We first performed normalization with the tools of `edgeR` package [47] for the RNA-seq data with only raw counts. Then, the samples were checked for genomic gene expression similarity and samples with inconsistent location in hierarchical clustering and principal component analysis plots, were treated as outliers and removed.

Next, the expression data, including both AD and normal samples, from different projects were treated as different batches and adjusted to remove the batch effects using `ComBat` [48]. The adjusted expression data were further normalized with quantile normalization and evaluated by PCA plots to make sure that the selected samples to have consistent expression profiles and have no clear batch effects among the data from different projects. Then, the resulting expression data are divided into two expression profiles for AD and normal samples, respectively.

The mouse and human brain microarray data were collected from `GEO` database. To minimize the effects of the different microarray platforms, we only selected the data performed with Affymetrix’s platforms. Normalized expression data were downloaded and used. For each dataset, we performed quality control to make sure the expression profiles of samples had good expression profile consistency with experimental descriptions introduced in the original paper. The batch effects of the data from different experiments were estimated and removed with `ComBat`. And then the data were combined together. They were further evaluated for expression profiles consistency and the experiment or sample outliers were removed. Finally, the probes of microarry were mapped to gene symbols. For the genes with multiple probes, we selected the ones with maximum expression values. The gene from human and mouse were mapped based on the gene homologous annotation from Mouse Genome Informatics (MGI)(www.informatics.jax.org).

### Differential co-expression analysis

Using the expression values of different samples as elements of expression vectors, the Spearman’s correlation of all gene pairs were calculated for AD and normal samples, respectively. To evaluate the robustness of calculated correlations, we performed two simulation studies. In the first evaluation, we randomly selected half of the samples and calculated new correlations. We then checked the mean and variance of correlation values under 100 rounds of simulation. The results helped us to understand the stability of observed correlation values under the different sample selection. In another evaluation, the annotation of samples for each gene was shuffled so that the gene pairs had the wrong sample mapping. The new correlations were calculated under 100 rounds of simulations. This simulation help us to evaluate the confidence ranges for the observed correlation values.

The correlation differences under disease and normal status were evaluated using the *R* package `DiffCorr` [24]. In this step, the correlation values were transformed with Fisher’s transform and z-scores were calculated to indicate the correlation differences. The p-value are calculated by fitting to a Gaussian Distribution. After comparing the efficiency, Benjamini & Yekutieli’s algorithm in *R* package [49] was implemented to control the false discovery ratio. At a cutoff of adjusted *p <* 0.01, we select the significantly differentially correlated genes.

### Enrichment analysis

The gene ontology (GO) annotation of gene lists was performed with the GO enrichment analysis tool `David` [37] under the default setting. The significantly enriched terms for biological process and cellular components were selected at a cutoff of *p* < 0.01. When multiple gene lists are available, the GO annotation results are visualized in a heatmap to facilitate comparisons.

In this work, we also performed enrichment analysis using an annotated gene list, e.g. DEGs and text-mining annotated genes. For *k* input genes, the number of genes with annotation is *x*. For *n* whole genomic genes, the number of genes with annotation is *p*. We use Fisher’s exact test to evaluate if the observed *x* genes result from random occurrences. We use the following R codes to calculate the p-value:

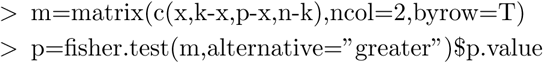

### Differentially expressed genes in Alzheimer’s disease

We performed differential expression analysis to the RNA-seq data collected in above steps from the AMP-AD program. For data from the MSBB project, four brain regions were treated as four independent datasets. Among 7 used datasets, some RNA-seq data had raw counts, e.g. data from MSBB project and we carried out differential expression analysis using `edgeR`. Other data with normalized expression values were analyzed by log2 transformed t-test. For all the datasets, the DEGs were determined at a cutoff of *p* < 0.01. To increase the confidences, the DEGs from different datasets were cross-validated with each other and the ones with clear inconsistency, e.g. the DEG list with weak overlap with other DEG lists were filtered out. Then, the selected DEGs were combined together as the DEGs of AD. The differential expression direction were also checked and determined by using the direction supported by the maximum datasets.

### Alzheimer’s disease related genes

The AD related genes were determined by selecting the ones with reported association with AD in published works, such as the genetics evidences and expression evidences. In this step, we use the gene-disease annotation for AD from IPA (http://www.ingenuity.com/products/ipa), Metacore (https://portal.genego.com/) and DisGeNet (www.disgenet.org/). We filtered out the low-confidence genes by manually removing the ones with only evidence of expression or those with inconsistency evidences.

### Gene co-expression network analysis

Gene co-expression network analysis was to find the gene clusters or modules with good co-expression similarity. As one of well recongnised implementation, `WGCNA` was applied to RNA-seq expression data following the protocol provided by the tool developers https://labs.genetics.ucla.edu/horvath/CoexpressionNetwork/Rpackages/WGCNA/[44]. All the expressed genes were clustered into modules, which were labeled with different colors. In each module, the connectivity and module membership of each module gene were calculated to assess the association and importance of genes in the modules.

## Results

### Transcriptional dysregulation in Alzheimer’s diseases

To find the dysregulation associated with AD, we evaluated the co-expression changes in the brain of AD patients, which indicate the transcription regulation changes. The whole process is detailed in Figure 1. In the first step (a), four sets of independent human RNA-seq expression data are collected from the `AMP-AD` program compiling with the data access control policy. The selected samples from different projects were processed and combined together to define the expression profiles of both AD and control subjects (see step (b)). Based on the clinical annotation provided by the data suppliers, 1045 AD samples and 622 control samples are determined and selected (see step (c)). As showed in Additional file 1, these samples have good homogeneity in their transcriptomic expression profiles and there is no clear outliers or batch effects. The mouse expression data were also collected from the `GEO` database (http://www.ncbi.nlm.nih.gov/geo/), where 931 samples from 20 microarray experiments are selected and processed to construct the mouse brain expression profiles.

**Figure 1.**
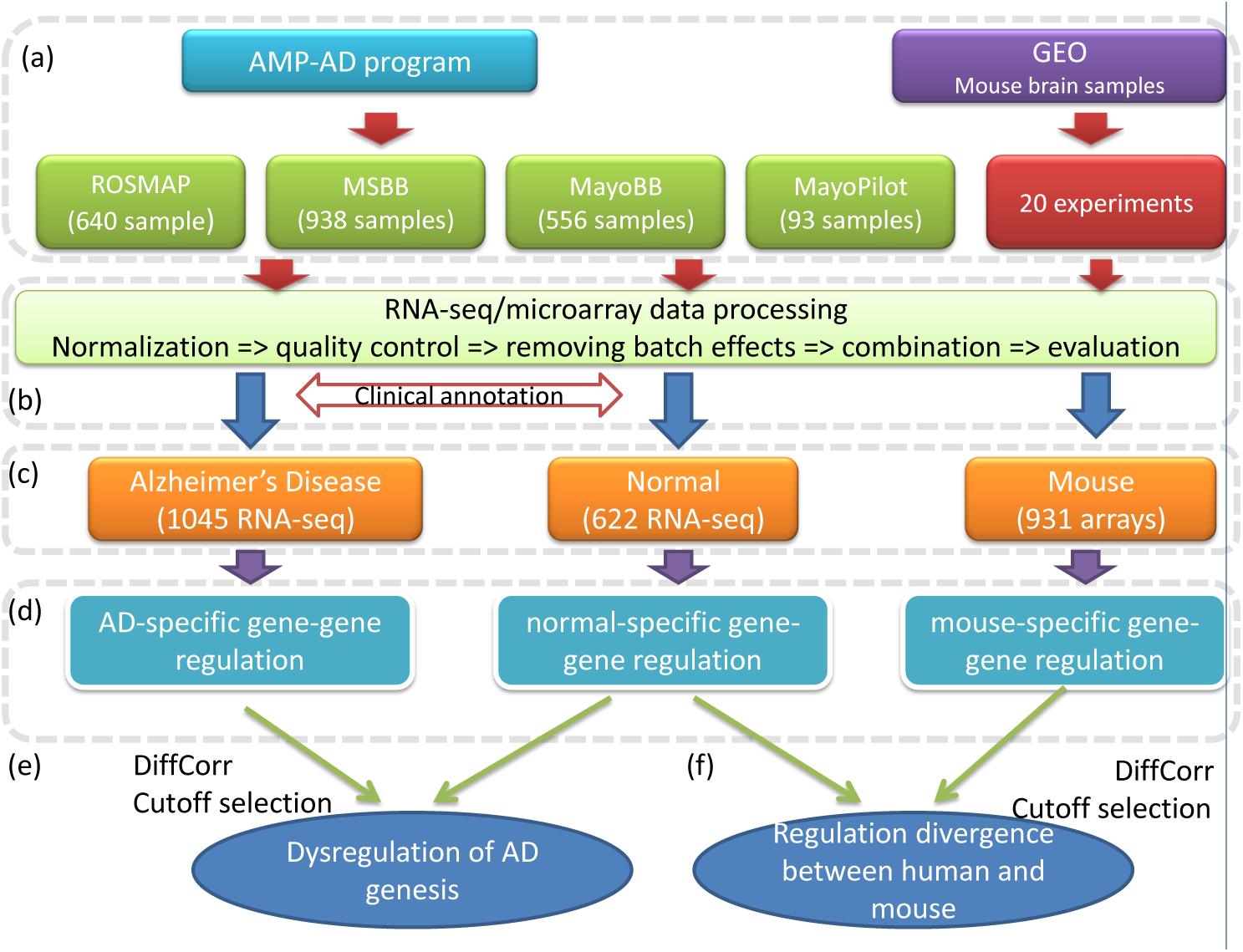
The pipeline to find the transcriptional dysregulation in AD. (a) The human RNA-seq expression data are collected from four projects of AMP-AD program; the mouse brain expression data are collected from 20 microarray experiments in `GEO` database. (b) The expression data from different datasets are processed, normalized and combined to describe the expression profiles of AD patients, control subjects and mouse. (c) The human brain samples are categorized into AD patients and normal samples based on the clinical annotation. (d) All gene pairs are evaluated for gene-gene regulations by measuring co-expression correlations for all the gene pairs. (e) The dysregulated genes are predicted by differential co-expression analysis. (f) The co-expression conservation of all gene pairs are compared between human and mouse

In the next step (d), the co-expression level, described by Spearman’s correlations, of all gene pairs are calculated for AD and control subjects, respectively. To check the robustness of calculated correlations, we performed random sampling and shuffling to original expression data. We found that the calculated values were tolerant to the sample selection and were less likely to result from random correlation (see Additional file 2 and Methods). These results suggest the correlation value to be robust enough to study the co-expression changes. In Figure 3(a), we show the plot for the correlation values of all gene pairs and find that the AD and normal samples have consistent co-expression profiles (Spearman’s *r* = 0.93). Applying the differential correlation analysis method introduced in [24], we found 87,539 out of 163 million gene pairs to have significant co-expression changes in the AD patients at a cutoff of adjusted *p <* 0.01 (see Additional file 3). Among them, there are 9168 genes with at least one DCE partner (see Additional file 4). Considering the nature of DCE analysis and the strict cutoff, all of the predicted gene pairs are co-expressed in either AD patients or normal people or both at a cutoff of *p* < 0.01 (see Additional file 3), which confirms that all the dysregulated gene pairs are potentially associated with the transcription regulation or co-regulation in the AD or normal status. Similarly, the DCE pairs are supposed to have altered regulation in the AD patients. Therefore, we can call them dysregulated genes.

We checked the co-expression status of the 87,539 dysregulated gene pairs. 22,150 gene pairs (25.3%) are co-expressed only in AD samples while 10,707 pairs (12.2%) are co-expressed only in normal samples at *p* < 0.01. Other gene pairs are coexpressed in both AD and normal samples. Among them, 31,685 pairs (36.2%) have increased co-expression correlation in AD while only 9694 pairs (11.1%) have decreased values. There are also gene pairs with reversed co-expression trend. For example, 6459 negatively correlated gene pairs (7.4%) in normal subjects become positive correlated in AD patients. Vice versa, 6844 gene pairs (7.8%) have the opposite changes. By summarizing the overall changes, we observed more gene pairs to have increased co-expression correlation in AD (2.6 times that of the gene pairs with decreased co-expression correlation), which suggests strengthened transcriptional regulation in the AD patients.

### Dysregulated AD genes

In published works, at least `777 expressed` genes have been reported to be associated with AD. We found that 479 of them were predicted with at least one dysregulated partner (see Additional file 5)(*p* = 2.5*e* − 9). Among them, the MAP1B gene was predicted with the maximum number of partners (365 genes), of which 32 genes were also AD associated genes. We found that the partners of MAP1B had diverse functions and many of them were associated with AD genesis, such as intracellular signaling cascades (42 genes, *p* = 3.7*e* − 4), regulation of apoptosis (18 genes, *p* = 3.4*e* − 3), neuron and dendrite development (5 genes, *p* = 4.13e — 3) and RNA metabolic process (19 genes, *p* = 4.71*e* − 3). This result confirms MAP1B, as a microtubule associated protein, to have diverse involvement in neuron related biological processes [25, 26, 27, 28] and suggests its important roles in the AD genesis [29]. Another example is the TREM2 gene, which is supported to be involved in the neuroimmunology of AD [30]. We observed 4 partners, including SLA (DCE at *p* = 3.7*e* − 10), HCLS1 (*p* = 2.5*e* − 10), C3AR1 (*p* = 8.7*e* − 10) and FCER1G (*p* = 1.3*e* − 9), all of which are associated with inflammation related functions. Among the well studied AD drug targets [31], we find their dysregulation, such as the amyloid precursor protein gene (APP, 136 partners), glycogen synthase kinase 3 beta (GSK3B, 34 partners) and BACE1 (8 partners), suggesting their transcriptional involvement in the genesis of AD.

In Table 1, we show 68 dysregulated AD genes, which are selected based on their partner numbers. To study their biological involvement, we studies the enrichment of AD related genes in their partners. We found the partners of 48 dysregulated AD genes not to be enriched with AD genes at a cutoff of *p* < 0.05, which suggested that these dysregulated AD genes were involved in AD genesis by affecting the genes without clear reports for AD association. We also checked the transcriptional association of 68 dysregulated AD genes. We found 56 genes to be differentially expressed (*p* = 0.05) in the AD patients and the significance for such enrichment was *p* = 1.1*e* − 27, which suggested the dysregulated AD genes to be associated with the transcriptional dysregulation even though most of the AD genes are not identified by differential expression analysis in published works. Similarly, we found the partners of 53 dysregulated AD genes to be enriched with the differentially expressed genes (DEGs), which confirms the transcriptional involvement of the dysregulated AD genes. Another investigation was to the aging related genes based on the annotation in our previous work [32]. We found 21 out of 68 dysregulated AD genes to be aging genes and the significance for such enrichment was *p* = 6.9*e* − 5. The partner genes of 39 dysregulated AD genes were also enriched with the aging genes at a cutoff of *p* < 0.05, confirming the association of the aging process with the genesis of AD.

**Table 1.**
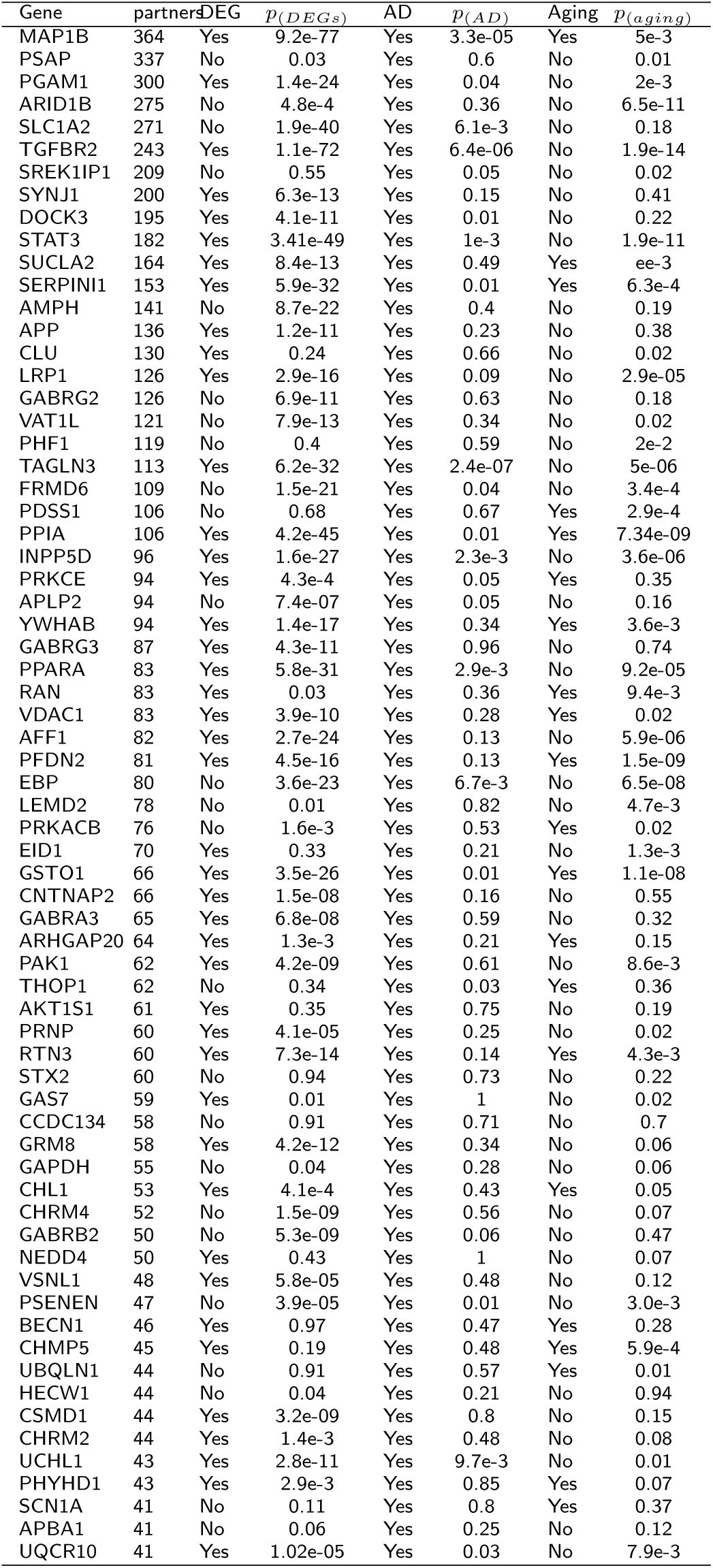
The dysregulated AD genes

We also studied the functional involvement of the dysregulated AD genes. In Figure 2, we show the functional annotation for the 68 dysregulated AD genes based on `Gene Ontology` annotation. Consistent with the selection criteria for AD genes, we found these genes to be associated with many AD related functions. Among them, “transmission of nerve impulse” is predicted to be the most enriched term (12 genes, *p* = 2.3*e* − 7). In the published works, synaptic dysfunction has been widely reported for its association with AD [33, 34] and our analysis confirms it to be one of the most affected processes. Other AD related terms include “neurological system process” (14 gene, *p* = 1.54*e* − 3), “regulation of protein kinase activity” (7 genes, *p* = 3.5*e* − 3).

**Figure 2.**
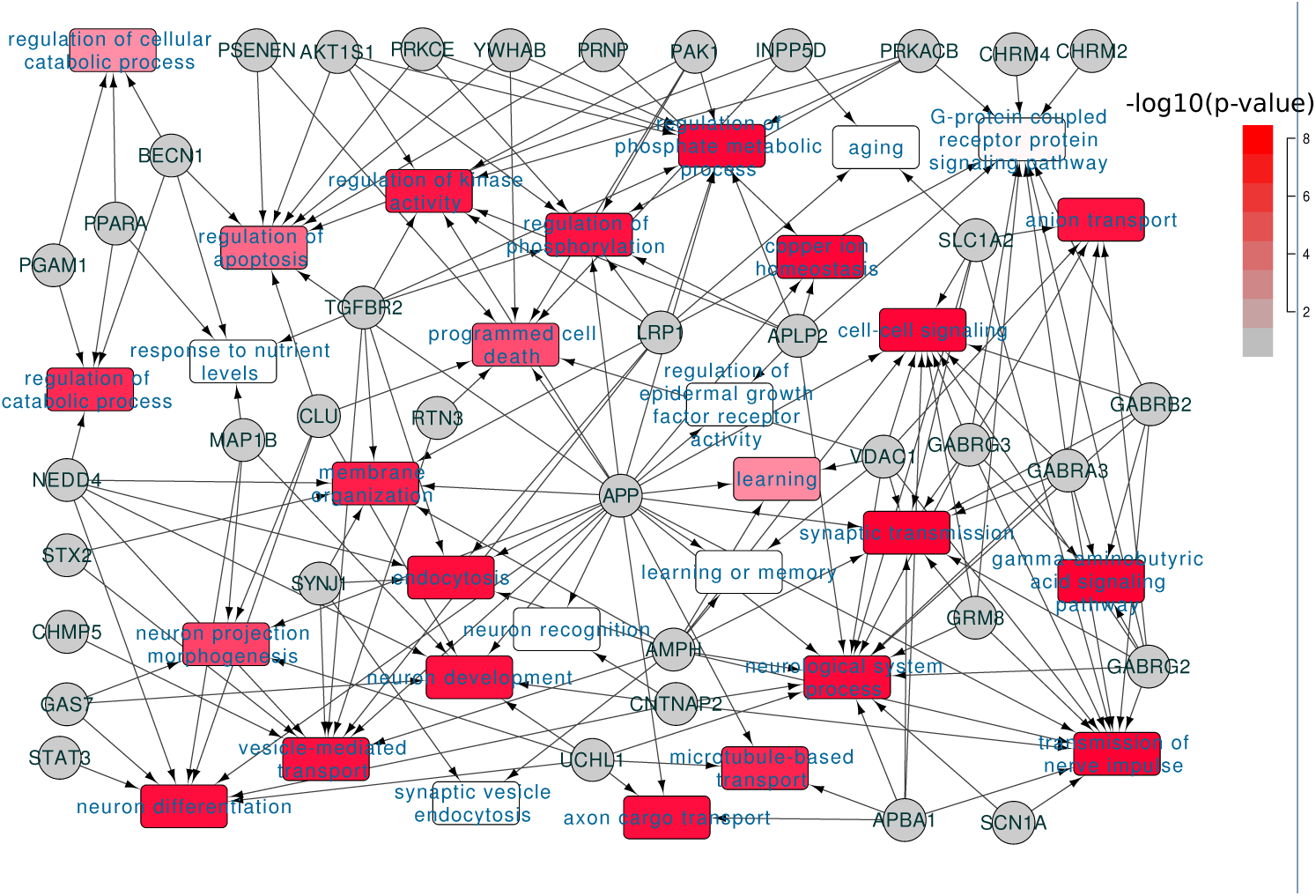
Functional invovlement of dysregulated AD genes. 6B dysregulated AD genes were annotated for their functional involvement based on the Gene `Ontology` annotation; the colors of GO terms indicated the enrichment significance.

In summary, we found the subset of the AD related genes that were transcriptionally dysregulated in AD; these genes were involved the genesis of AD by affecting the AD related pathways or biological processes, e.g. synaptic transmission.

### A core network involves the dysregulation of AD

Following the widely accepted hypothesis that the hub genes of a network take more essential roles, we assume that the genes with more dysregulated partners will have more essential roles in the etiology of AD [35]. Another assumption is that the dysregulated genes and their partners would participate in the same pathways or biological progresses. Therefore, we can infer the function of dysregulated genes by analysis of their partners [36].

Based on these assumptions, we defined the genes with both transcriptional dysregulation and involvement in the genesis of AD by the following criteria: (1) with more than 50 partners; and (2) to be differentially expressed in AD (*p* < 0.01, see Additional file 6), which ensures the selected genes to be transcriptionally associated with AD, and their partners to be enriched with AD genes; or (3) reported as AD associated genes and their partners enriched with the differential expressed genes at a cutoff of *p* < 0.01. In this way, 97 dysregulated genes are selected (see Figure 1(c, d) and Additional file 4). Among them, 64 genes are differentially expressed in AD and 39 genes are reported as AD associated genes. These genes are dysregulated in 14,322 gene pairs with 3681 partners. Of the dysregulated genes, TMEM178A is the most dysregulated gene with 669 partners. We checked the co-expression status between dysregulated genes and their partners and found 85.9% of the gene pairs to have increased co-expression correlation values, which was far more than the observed percentage (66.2%) with non-filtered dysregulated gene pairs. In AD patients, we also observed 3461 gene pairs to be co-expressed only in AD, which was 3.6 times the normal-specific co-expressed pairs (*p* = 4.6*e* − 67), suggesting that the AD patients have more and strengthened transcriptional regulations.

Even though the dysregulated genes were not selected for any co-expression correlation among each other, we observed the 97 dysregulated genes to have stronger co-expression correlation values than random selected genes (see Figure 3(b)). Of 4656 gene pair combinations, 95.7% of them are co-expressed at a cutoff of *p* < 0.01 and 86.9% have a correlation *r* > 0.3. The median correlation value is 0.556 (*p* = 7.7*e* − 86). We also checked their co-expression changes and found 929 out of 4656 gene pairs to be differentially co-expressed in the transition from normal to AD.

**Figure 3.**
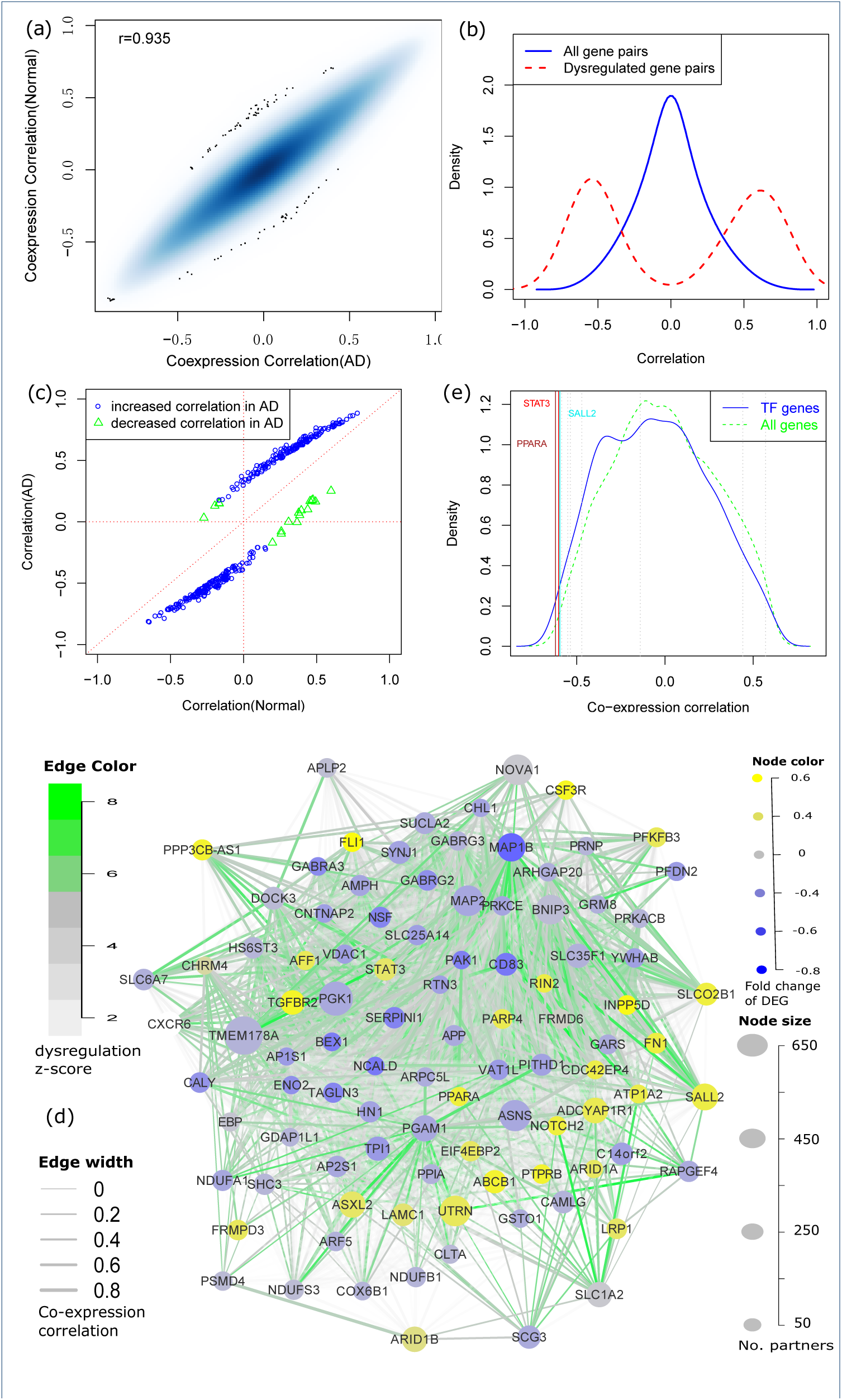
A core network involving the dysregulation of AD. (a) The co-expression profiles of the genomic genes are highly similar between human AD and normal samples (*r* = 0.935); (b) The gene co-expression correlations among dysregulated genes (red dash line) and genomic genes (blue solid line), which indicates that most of the dysregulated genes interact with each other; (c) Majority of the dysregulated genes have increased correlations (both direction) in the transmit from normal to AD; (d) 96 (out of 97) dysregulated genes can be interconnected as a regulatory network based on co-expression and differential co-expression information. (e) TMEM178A expression is negatively correlated with dysregulated transcription factors, among which PPARA is ranked as the most negatively correlated transcription factors.

Using the co-expression and differential co-expression information, we can connect the dysregulated genes as a network (see Figure 3(d)). In this network, 96 out of 97 nodes can have at least one connected co-expression partner at *r* > 0.3 and 90 nodes can also have at least one dysregulation partner at a cutoff of adjusted *p <* 0.01. Combining the co-expression and differential co-expression information, 96 out of 97 dysregulated genes are connected with other dysregulated genes. The island of this network is ARF5, which is differentially expressed in AD but not reported with any association with AD. Therefore, we filtered it in Figure 3(d). We further checked the direction of edges and found 870 out of 929 dysregulated edges to have increased co-expression correlation in AD (see Figure 3(c)) while 176 of these regulations were AD-specific. In Figure 3(d), we also show the differential expression information of 96 nodes and find 26 up-regulated and 37 down-regulated genes in AD. The transcriptional regulations are observed between up- and down-regulated genes, including 42.7% of co-expressed edges to have negative correlation values.

In the network, we observed 10 transcription factors (TFs): FLI1, NOTCH2, SALL2, STAT3, PPARA, BEX1, ARID1A, ARID1B, AFF1 and PRNP. We checked if these 10 TFs regulated the expression of other dysregulated genes. Due to limitation for experimental validation, e.g. human brain sample collection and manipulation, we evaluated their regulatory roles by comparing their expression profiles with the dysregulated genes. Compared with 2405 annotated TF genes in the `Gene Ontology`, we observed that these TF genes were always ranked as the most coexpressed TF genes. In Figure 3(d), we show the example for TMEM178A gene. We found 9 TFs to have strong co-expression correlations, especially for PPARA and STAT3, which were co-expressed with TMEM178A at *r* = −0.62 and *r* = −0.603, ranked as the 10th and 16th of the most negatively co-expressed TF genes. Similar results were observed with other dysregulated genes (see Additional file 7). Another investigation we carried out was to predict the gene-specific regulators using the method introduced in [36]. In this step, we predicted the enriched regulators for each of 97 dysregulated genes using AD and normal expression data as the input expression matrix. In the Jaspar database, only STAT3 has clear TF binding profiles and therefore, we can only predict the transcriptional regulation for STAT3. We found that STAT3 was ranked as the 16th of the most important regulators for 96 dysregulated genes in 474 annotated TFs. In normal and AD samples, STAT3 was predicted to regulate 25 and 31 dysregulated genes, respectively. Overall, these results suggest that the dysregulated TFs may be involved in transcriptional regulation or dysregulation in AD.

### Aging, synaptic transmission and metabolism are dysregulated

Applying a strategy of `guilt` by `association`, we extended the functional study of 97 dysregulated genes to the subnetworks comprising of themselves and their partners. In this step, we used each of 97 dysregulated genes as hub and exacted the partner genes to construct the dysregulation subnetwork, which could be treated as the co-regulated unit for further studies. Investigation of these subnetworks suggested them to have overall consistent co-expression changes. Taking the TMEM178A subnetwork as an example, we found 665 out of 669 nodes to have increased co-expression correlations with the hub gene, including 164 gained coexpression in AD. Similar results can be observed with many other dysregulated genes (see Additional file 4).

We investigated the association of subnetworks with AD genesis. The first attempt was to check the enrichment of DEGs in AD. We found all the subnetworks (97/97) to be enriched with DEGs at a cutoff of *p* < 0.01 (see Figure 4(a)). Taking the TMEM178A subnetwork as an example, 413 out of 669 partner genes was differentially expressed in AD and the significance for this enrichment is *p* = 6.5*e* − 127 by Fisher’s exact test. By checking the other genes, we found 84 subnetworks to be enriched with DEGs with a high statistical significance of less than *p* = 1*e* – 10. Such significant enrichment suggests all the subnetworks to be associated with the genesis of AD at the transcriptional level.

**Figure 4.**
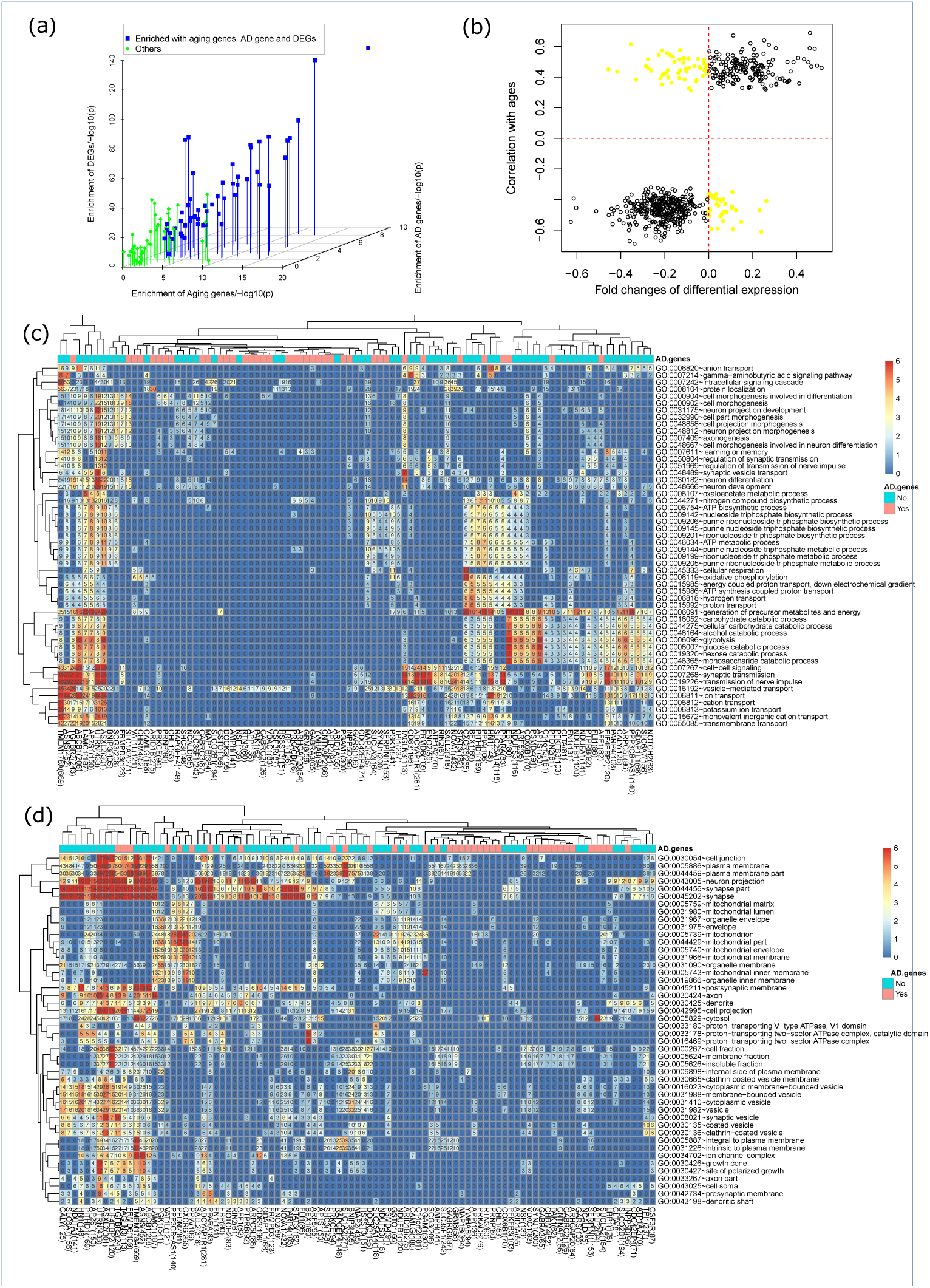
Aging, synaptic transmission and metabolism are dysregulated. (a) The enrichment of aging genes, AD differentially expressed genes and AD related genes in 97 subnetworks; (b) the dysregulated genes usually have the same expression change direction in the aging process and the AD genesis; (b) functional enrichment analysis to dysregulated genes for (c) biological process and (d) cellular component.

Considering the fact that age is the biggest risk factor for AD, we checked the enrichment of aging related genes in each subnetwork which were identified by finding 2862 genes with aging associated expression or DNA methylation in our previous work [32]. For the subnetworks with differentially expressed genes as hubs, we found 53 out of 64 subnetworks to be significantly enriched with aging related genes (see Figure 4(a)). Taking the TMEM178A subnetwork as an example, we found 165 out of 669 genes to be aging related genes (*p* = 3.2*e* − 20). Out of 2862 aging genes, 624 genes are also dysregulated in AD. We further evaluated the involvement of dysregulated aging genes in the genesis of AD by checking their differential expression direction in AD and their expression trend in the aging process. As showed in Figure 4(b) 541 dysregulated aging genes have the same expression direction, e.g. the aging related genes with increasing expression levels in the aging process would have up-regulated expression in the AD patients and vice versa. Overall, the analysis results suggest that the aging genes will be associated with the transcriptional dysregulation of the AD genesis.

We performed functional annotation using `David` [37] for each subnetwork. Figure 4(c) shows the enriched biological processes (BPs). The most enriched category is for the synaptic function related terms. One example is the “transmission of nerve impulse” term, which is enriched in 37 subnetworks and that is also the most enriched term for most of the 37 subnetwork. The other related terms include “synaptic transmission” and “cellcell signaling”. In published reports, these BPs have been reported for their involvement in the AD genesis [33, 34]. Another category is the metabolism related terms. Among them, “generation of precursor metabolites and energy” was significantly enriched in 39 subnetworks and “glycolysis” was significantly enriched in 29 subnetworks. The links between metabolism and AD are also supported by the published works [38, 39]. Even as different functional categories, synaptic function and metabolism related terms are usually enriched by the same subnetworks. We also observed other enriched terms and most of them have been reported for association with AD, such as “cation transport” [40], “ATP metabolic process” [41], “learning or memory” [41] and “neuron differentiation”. Using `David`, we also checked the cellular component (CC) enrichment (see Figure 4(d)). We found neuron, especially synapse related CCs to be the most enriched cellular location. Consistent with BP analysis results, the “synapse part” term was enriched in 46 subnetworks, especially the subnetwork with dysregulated DEGs as hubs, which confirmed the analysis results with BP terms.

As described above, the dysregulated genes are either DEGs or literature reported AD genes. We found the functional involvement for the two groups of subnetworks to be different. The 39 subnetworks with hubs of dysregulated AD genes are less associated with any enriched biological process. For example, the term “transmission of nerve impulse” is enriched in only 8 out of 39 subnetworks while there are 29 out of 64 DEGs subnetworks to be enriched with this term. Similarly, the other AD associated terms are also less likely to be enriched with these 39 subnetworks. However, considering the fact that the dysregulated AD genes are significantly associated with the AD related terms (see above section), including synaptic function and metabolism related biological processes, we can assume that the 39 subnetworks with the hubs of dysregulated AD genes are also associated with the enriched terms.

### Association with clinical outcomes

In the `ROSMAP` project of `AMP-AD` program, most samples have been annotated with clinical information. We studied the association of gene expression with three disease relevant clinical traits, e.g. cognitive test scores (cts), braak stage (braaksc) and assessment of neuritic plaques (ceradsc). We did not find any gene to have strong expression correlations (e.g. *r* > 0.6) with studied traits. The maximum correlation was observed with SLC6A9 at *r* = −0.388. For all the genes, only 12.3% of them have a clinical association at |*r*| > 0.15. This result suggests that the clinical outcomes of AD may be affected by combined effects and even beyond the effects of gene expression.

In Figure 5, we show the clinical association results for 97 dysregulated genes. We found that the dysregulated genes were always ranked as the most clinical associated genes. Of 97 dysregulated genes, 66 genes have a clinical association of |*r*| > 0.15 (*p* = 4.49*e* − 37). Among them, NCALD, a neuronal calcium binding protein, was associated with the cognition score at *r* = 0.268.

**Figure 5.**
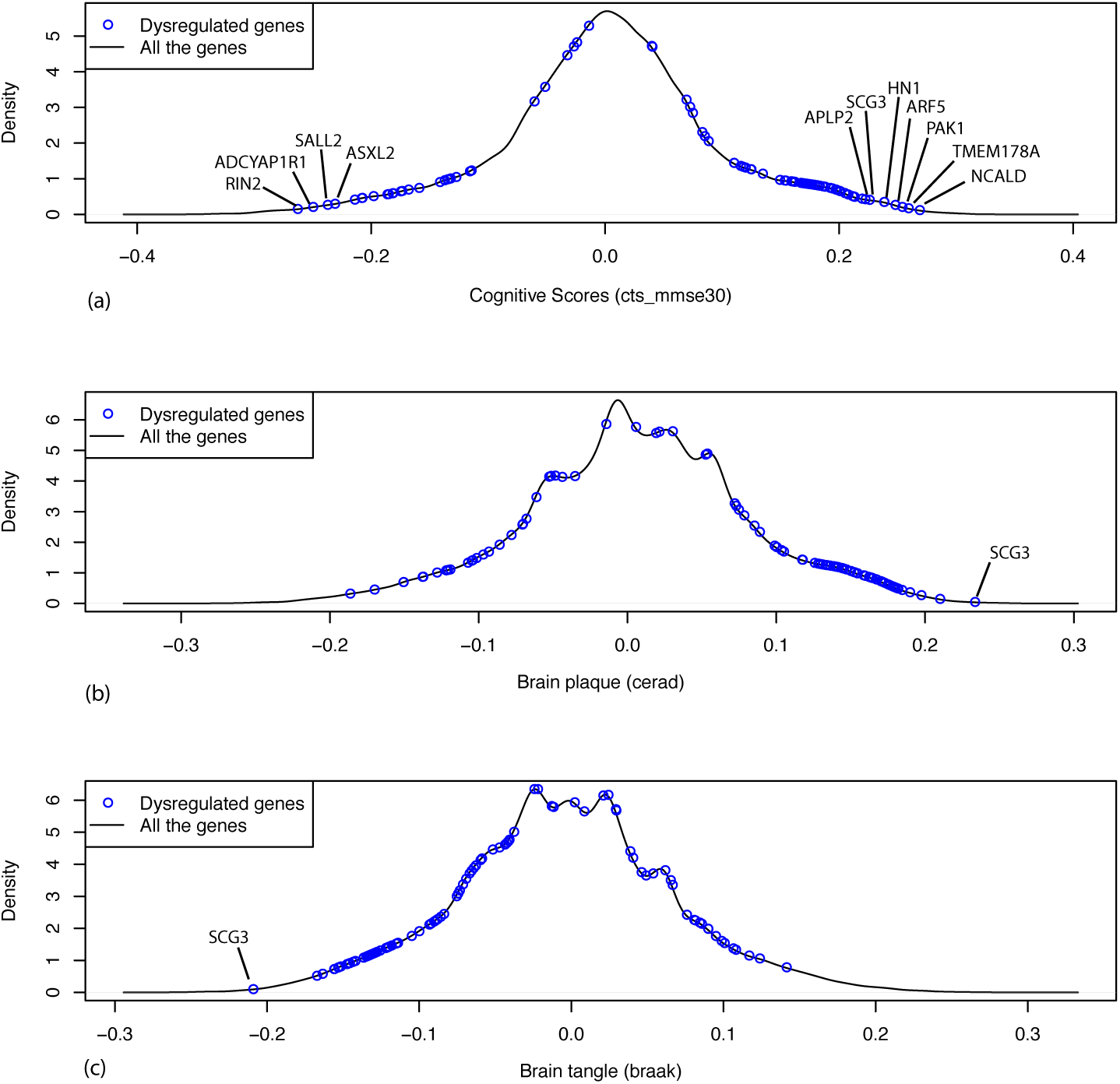
The dysregulated genes prefer to have more association with clinical traits, including (a) cognitive test score (cts), (b) braak stage (braaksc) and (c) assessment of neuritic plaques (ceradsc). The black lines show the density plots of clinical association (correlation) for all protein coding genes; the blue points indicate the 97 dysregulated genes.

Another investigation is to the combined effects of dysregulated gene pairs. We studied the partial correlations between the dysregulated genes and the clinical traits by controlling the effects of their dysregulated partners. We found that many dysregulated gene pairs had improved clinical association (see Table 2 and Additional file 8). In these dysregulated gene pairs, the clinical association of one gene is improved by considering the gene expression of its partners, which indicates the importance of dysregulation relationship.

**Table 2.** The association of dysregulated genes with the cognitive test scores

| gene1 | gene2 | coexpression cor. |  | cor. with cts<sup>1</sup> |  | partial cor.<sup>2</sup> |  |
| --- | --- | --- | --- | --- | --- | --- | --- |
|  |  | ad | normal | gene1 | gene2 | gene1 | gene2 |
| RPS19 | APLP2 | -0.446 | -0.696 | -0.036 | 0.226 | 0.205 | 0.299 |
| SLC35F1 | PPP2CA | 0.718 | 0.528 | 0.039 | 0.208 | -0.183 | 0.271 |
| SLC35F1 | RTN3 | 0.711 | 0.517 | 0.039 | 0.217 | -0.180 | 0.273 |
| RALGPS1 | APLP2 | -0.039 | 0.265 | 0.030 | 0.226 | -0.188 | 0.289 |
| SLC1A2 | FAT1 | 0.233 | 0.511 | -0.052 | -0.185 | 0.153 | -0.232 |
| SLC15A2 | SLC1A2 | 0.442 | 0.693 | -0.208 | -0.053 | -0.298 | 0.214 |
| SDC2 | SLC1A2 | 0.359 | 0.611 | -0.199 | -0.053 | -0.268 | 0.180 |
| NOTCH2 | SLC1A2 | 0.172 | 0.533 | -0.210 | -0.053 | -0.267 | 0.177 |
| SYNJ1 | PRUNE2 | 0.333 | 0.581 | 0.160 | 0.062 | 0.218 | -0.163 |
| GPRC5B | SLC1A2 | -0.055 | 0.27 | -0.276 | -0.053 | -0.316 | 0.169 |
| SUCLA2 | PRUNE2 | 0.311 | 0.557 | 0.153 | 0.062 | 0.210 | -0.158 |
| SUCLA2 | ARFGEF2 | 0.746 | 0.861 | 0.153 | 0.067 | 0.201 | -0.148 |
| PRKACB | CDS2 | 0.641 | 0.794 | 0.187 | 0.070 | 0.225 | -0.145 |
| PRKCE | KCNA1 | 0.62 | 0.397 | 0.196 | 0.062 | 0.237 | -0.148 |
| APP | RBMX2 | -0.037 | -0.353 | 0.169 | -0.064 | 0.195 | 0.118 |

<sup>1</sup> the correlation between gene expression and cognitive test score<sup>2</sup> the partial correlation with cognitive test score

### Dysregulation divergence in mouse

Mice are widely used as a model animal in wet-lab studies for AD. Investigation of the evolutionary conservation of gene regulation can help with the evaluation of experimental studies drawn in mice. Therefore, we collected mouse brain microarray expression data from `GEO` database. 1583 samples from 24 experiments were identified (see Additional file 9). We removed arrays with poor quality or inconsistent expression profiles and finally, 931 samples from 20 experiments were used. The selected samples were combined together to describe the mouse brain expression profiles. Using gene homologous annotation from Mouse Genome Informatics (MGI) (www.informatics.jax.org), 14186 expressed genes were used for co-expression evaluation with the human normal control samples. We first studied the overall gene expression similarity between mouse and human. As showed in Figure 6(a), we found human and mouse to have consistent expression profiles with the median Spearman’s correlation among samples at *r* = 0.69, which indicated the conservation of gene expression profiles between human and mouse.

**Figure 6.**
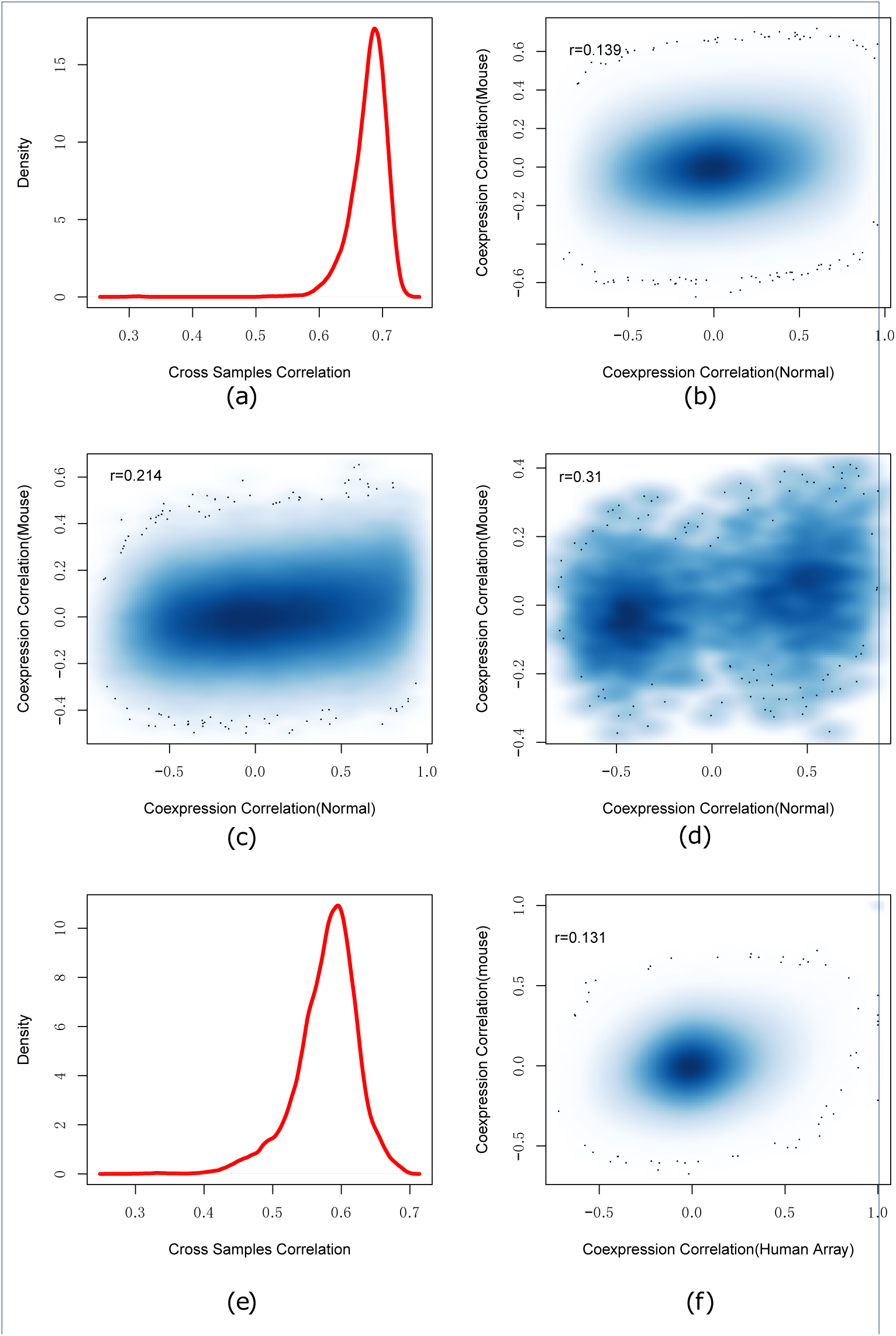
The dysregulation divergence between human and mouse. (a) shows the density plot of expression profile similarity (correlation) of human and mouse homologous genes, which suggests the conserved gene expression patterns between human and mouse. However, (b) the regulations among all the genomic genes are not conserved between human and mouse (*r =* 0.139). By restricting the studied regulations to the dysregulated pairs (c) and the edges in Figure 3(d), the conservation between human and mouse regulations was weakly improved. The observation was further investigated with human microarray data and found good consistent with human RNA-seq (e) but still failed to suggest any gene-gene regulation conservation between human and mouse (f).

Another investigation was to co-expression of gene pairs. The co-expression correlations of all gene pair combinations were calculated for mouse and human normal samples, respectively. Figure 6(b) showed the co-expression correlation values. Human and mouse had different co-expression profiles and the Spearman’s correlation to co-expression correlations was only *r* = 0.139. Further, we performed differential co-expression analysis and found 25.0% of gene pairs to have differential coexpression at the cutoff of adjusted *p* < 0.01. We also checked the co-expression directions and found 34.7% of 10 million co-expressed gene pairs to have inconsistent correlation directions, which indicated the divergence of co-expression patterns between human and mouse. Similar results were also observed with the human AD samples (see Additional file 10). To understand the co-expression conservation of AD related genes, we restricted the same analysis to the 87539 differential correlated gene pairs. As shown in Figure 6(c), improved conservation was observed (*r* = 0.214, *p* = 6.46*e* − 266). We further restricted the analysis to 97 dysregulated genes and found the co-expression conservation were improved further (*r* = 0.31, *p* = 8.08*e* − 6) (see Figure 6(d)).

Different platforms, e.g. RNA-seq and microarray, may lead to different expression measurements [42, 43]. Therefore, we constructed the human brain expression profiles by collecting the microarray data from GEO database (see Additional file 11). To minimize the influences of different platforms, we only collected data from AffyMetrix’s platform when the mouse expression data were also generated using AffyMatrix’s technology. Finally, 452 samples were selected to describe the human brain expression profiles. As showed in Figure 6(e), we found that the expression profiles measured by microarray and RNA-seq were not completely consistent(*r* = 0.54) (see Figure 6(e)). Next, we re-evaluated the co-expression conservation using the coexpression profiles calculated from human brain microarray data. The Spearman’s correlation to the co-expression correlations was *r* = 0.152 (see Figure 6(f)), which still indicated strong dysregulation divergence between human and mouse.

In summary, human and mouse may have the divergent co-expression patterns in brain, even though they have the conserved gene expression profiles, which indicates the different transcriptional regulation.

### Comparison with established works

Based on the assumption that the connectivity in the co-expression network indicates the gene importance, connectivity has widely been used to identify the essential components for a specific biological process. We investigated the association between dysregulation and the connectivity in the co-expression network. We firstly checked the connectivity preference of dysregulated genes by selecting the top 100 dysregulated genes discovered in the above analysis. As shown in Figure 7(a)), we failed to observe any connectivity preference when compared with genomic genes (*p* = 0.229). Another investigation was to check if the genes with stronger connectivity would be more differentially co-expressed. As shown in Figure 7(b) and Additional file 12, the maximum and median differential correlation did not show strong association with the connectivity, including the most dysregulated ones.

**Figure 7.**
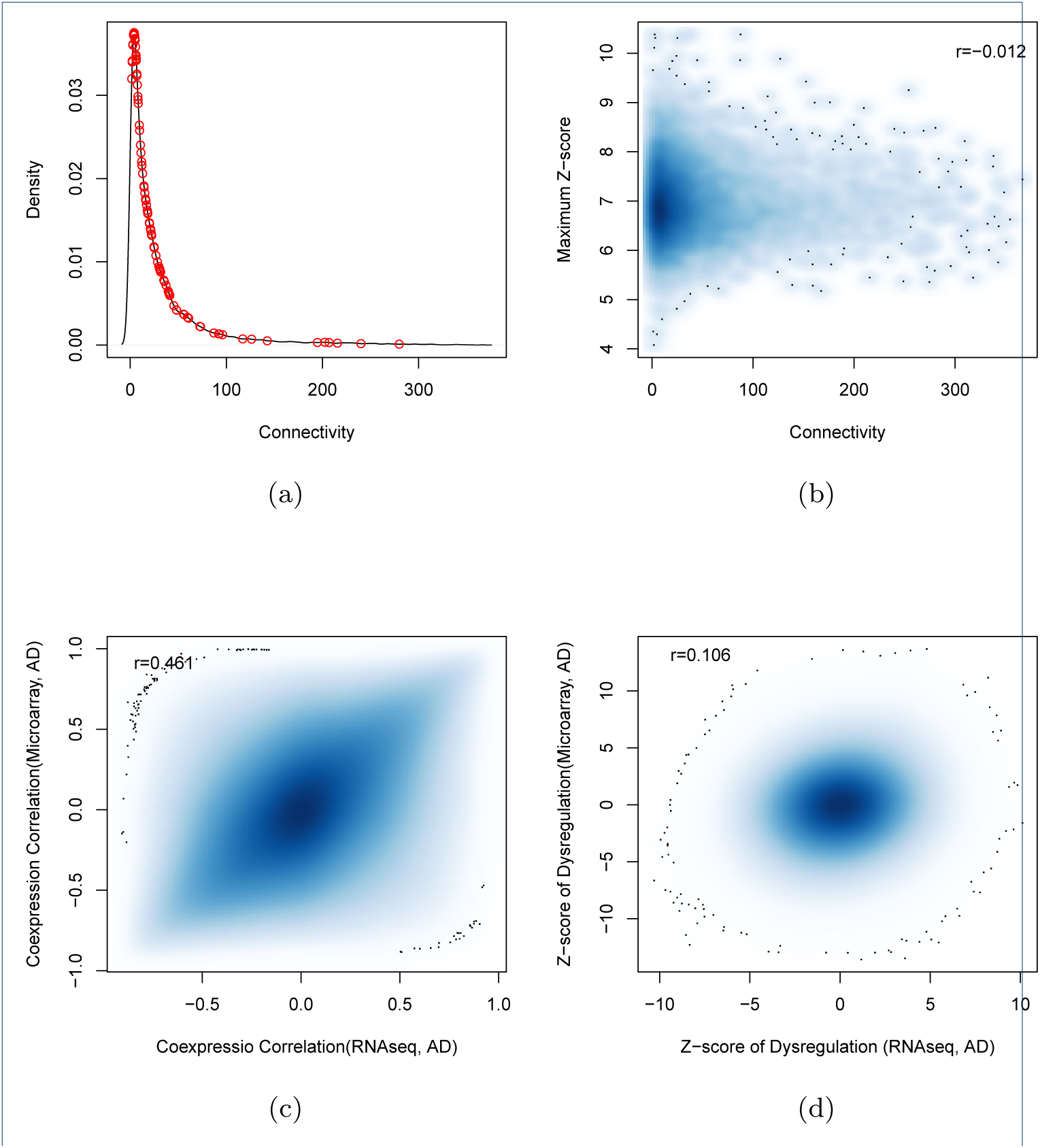
Comparison analysis with the results from estiblished work. (a) The connectivity distribution of top 100 of the most dysregulated genes (red circle points) fails to indicate any association between connectivity and dysregulation; (b) the same result is observed with all the genomic genes (*r* = −0.012). (c) using human RNA-seq and micorarray data, the calculated gene co-expression correlations are consistent; but (d) the predicted dysregulation are not consistent.

Next, we investigated if the dysregulated genes could be identified by co-expression network analysis. We applied `WGCNA` [44], a popular co-expression network analysis tool, to human brain RNA-seq expression data and 43 modules were predicted. The 97 dysregulated genes were mapped into 22 modules. We found the number of mapped dysregulated genes to a specific module was correlated with its module size (*r* = 0.76). For example, the BLUE module had the largest module size of 7776 genes and it was also mapped with the maximum number of dysregulated genes (44 genes) (see Additional file 13). Further, we ranked the module members based on their connectivity in the modules. None of 97 dysregulated gene was ranked as the most connected hubs. All these results suggest that the dysregulated genes are not necessary to be discovered by co-expression network analysis.

In independent work, the dysregulation of AD has been investigated using microarray data [22], where the dorsalateral perfrontal cortex region from 310 AD patients and 157 normal subjects was measured by microarray. Considering the fact that platforms may affect the analysis results [43], we compared their analysis results using 13606 common expressed genes. The gene pairs from microarray and RNA-seq platforms had overall consistent co-expression profiles (*r* = 0.49, *p* = 1.2*e* − 199) (see Figure 7(c)). Applying the differential correlation test, we found 18.5% of gene pair combinations to be differentially co-expressed, which was far more than the number of predicted dysregulations for AD genesis: that is, 0.05% of all gene pairs for RNA-seq dataset and 0.6% for microarray datasets. This result suggested the dysregulations predicted using microarray and RNA-seq data to be different. To illustrate it, we checked the consistency of predicted dysregulation by DCE analysis for both RNA-seq and microarray data. Figure 7(d) shows the dysregulation z-scores of all gene pairs, which describe the significance of co-expression changes. We found that the overall dysregulation were weakly consistent between RNA-seq and microarray (Spearman’s *r* = 0.106). At a cutoff of adjusted *p* < 0.01, 59,710 and 603,335 gene pairs were predicted to be dysregulated with RNA-seq and mi-croarry data, respectively. Among them, only 1469 gene pairs were shared by two datasets. We also checked the predicted partner number of dysregulated genes from RNA-seq and microarray data and found them to be weakly consistent (*r* = 0.218) (see Additional file 14).

## Discussion

Using the RNA-seq data, we studied the association between gene expression correlation and connectivity. We found that the genes with higher connectivity showed stronger co-expression correlations (see Additional file 15(a)). Functional annotation to top 200 of the most connected genes suggested these genes more associated with house-keeping related roles, such as protein location and protein transport (see Additional file 15(b). Considering the fact that AD is associated with loss of neuron related functions, we further checked the connectivity of 2155 brain tissue-specific genes [45]. Even though the brain-specific genes have stronger connectivity than the genomic genes (*p* = 7.53*e* − 56), we found that only 14.8% of the the brain-specific genes were ranked in the top 1000 of the most connected genes. Similar results were observed with the AD related genes. All these observations suggest that the genes associated with AD genesis are not required to have strong connectivity and this led us to extend the connectivity-based analysis to a less biased investigation.

Therefore, we evaluated the co-expression differences between AD and normal samples for all the gene pair combinations without considering gene connectivity. The genes with differential co-expressed profiles are believed to be dysregulated in the genesis of AD. Using a similar assumption, the genes with the most numbers of dysregulation pairs are supposed to take more essential roles. Finally, 97 dysregulated genes were predicted. In consistent with our hypothesis, the predicted genes failed to show strong connectivity in co-expression network analysis. This was further supported by mapping these genes into the `WGCNA` analyses results. These results suggest that this work reveals the etiology of AD in an independent and novel view.

We performed functional annotation on the 64 dysregulated DEGs and found them to take diverse functions. The limited gene number and the relative independence of dysregulated genes hindered the efficiency of functional annotation. We extended the functional studies of dysregulated genes to the investigation to their dysregulated partners. We observed 33 of them to be associated with synaptic transmission related functions while synaptic dysfunction is believed to be one of the key characteristics of AD [34]. Another evaluation suggested the dysregulated partners to be enriched with the aging related genes, which explained the effects of age on AD genesis. Overall, the analysis results suggested the predicted dsyregulation to be related to the AD genesis and it indicated the involvement of transcriptional regulation among many reported mechanisms.

We defined the transcriptional “dysregulation” by selecting gene pairs with changed co-expression between disease and normal samples. This can take into account different regulatory mechanisms. The gene pairs can be down- or up-stream members in regulatory pathways, i.e. either A regulates B or B regulates A. They can also be co-regulated genes by common upstream pathways. In the context of co-expression or differential co-expression, it is not easy to elucidate the exact information of their relationship. However, by checking the functional annotation to the hub genes and their partners, we always found consistent functional annotation. For example, EIF4EBP2 is reported as a translation initiation repressor and is involved in synaptic plasticity, learning and memory formation [46]. Functional enrichment analysis to its partners suggests them to be associated with functions related to synaptic transmission, which confirms the consistent functional involvement between hub genes and their partners.

In differential co-expression analysis, the analysis results are subtle and related to the batch or platform effects of the data from independent projects. We applied several steps to improve the confidence. The most critical one was to collect large-scale RNA-seq data from the AMP-AD program, which minimized the effects of different platforms and batches. Due to the large-scale RNA-seq data, we were able to perform deep evaluation of the quality of expression data by comparing the sample consistency.

Another novel finding is the divergent transcriptional regulations between human and mouse. As the widely used animal model, many neurological studies are carried out in mouse. Our finding put a potential warning to some wetlab conclusions especially these from transcriptional related studies.

## Acknowledgment to the Data Contributors

The results published here are in whole or in part based on data obtained from the Accelerating Medicines Partnership for Alzheimer’s Disease (AMP-AD) Target Discovery Consortium data portal and can be accessed at doi:10.7303/syn2580853.

The full acknowledgment to the individual data contributors are available in Additional file 16.

## Author’s contributions

GM conceived of the study, designed the study, carried out the analysis and drafted the manuscript. XG collected the data, participated in its design and coordination and helped to draft the manuscript. HM collected the data, conceived of the study and participated in its design and drafted the manuscript. All authors read and approved the final manuscript.

## Ethics approval and Consent to Publish

Not applicable

Consent for publication

Not applicable

## Acknowledgements

We thank Dr. Xiaoyuan Zhou and Yanjiao Zhou for reading manuscript and giving us many useful suggestions. We are grateful to Dr. David Tattersall for his review of the manuscript.

## Tables

### Additional Files

Additional file 1

Genomic gene expression profile homogeneity evaluation for the samples from independent RNA-seq projects. The sample distribution in the principal component analysis (PCA) plot indicates the expression similarity of the selected samples for AD (a) and normal samples (b), respectively.

Additional file 2

Evaluation of the co-expression correlations by randomly sampling (a) and shuffling (b). (a) Half of the sample are randomly selected to calculate the new correlations. (b) All the genes are shuffled with random samples annotation so that the gene pairs have the wrong sample mapping.

Additional file 3

The 87,539 dysregulated gene pairs between AD and normal

Additional file 4

All the dysregulated genes with at least one partner. Other annotation including differential expressed in AD samples, aging related genes, AD genes, connectivity in co-expression network.

Additional file 5

The Alzheimer’s disease related genes collected by text mining to the published works.

Additional file 6

The differentially expression genes in AD patients.

Additional file 7

The co-expression between four TF genes and dysregulated genes. The solid lines indicate the co-expression correlation distribution between dysregulated gene and 2045 TF genes; the dash line indicate the co-expression correlation between dysregulated genes and the genomic genes; the legend shows the TF genes with maximum co-expression correlation with dysregulated genes.

Additional file 8

The association between dysregulated genes and clinical traits.

Additional file 9

The used microarray data for mouse brain.

Additional file 10

Mouse may have divergent co-expression profile with human AD samples.

Additional file 11

The used microarray data for human brain.

Additional file 12

The association between dysregulation and connectivity in the co-expression network. The y-axis shows the median z-score of the differential co-expression.

Additional file 13

The dysregulated genes and the WGCNA analysis results.

Additional file 14

The partner number of dysregulated genes predicted using RNA-seq and microarray data.

Additional file 15

The association between connectivity and co-expression correlation. (a) the genes with higher connectivity are always the genes with higher co-expression correlation. (b) Functional annotation to the top 200 genes with highest connectivity.

Additional file 16

The full acknowledgment to the individual data contributors.

